# PTPN22 R620W Gene Editing in T Cells Enhances Low Avidity TCR Responses

**DOI:** 10.1101/2022.07.21.500924

**Authors:** Warren Anderson, Fariba Barahmand-pour-Whitman, Peter S Linsley, Karen Cerosaletti, Jane H Buckner, David J Rawlings

## Abstract

A genetic variant in the gene *PTPN22* (R620W, rs2476601) is strongly associated with increased risk for multiple autoimmune diseases and has been linked to altered TCR regulation. Here, we utilize Crispr/Cas9 gene editing with donor DNA repair templates in human cord blood-derived, naive T cells to generate *PTPN22* risk edited, non-risk edited (silent modification), or knock out T cells from the same donor. *PTPN22* risk edited cells exhibited increased activation marker expression following non-specific TCR engagement, findings that mimicked PTPN22 KO cells. Next, using lentiviral delivery of T1D patient-derived islet-antigen specific TCRs, we demonstrate that loss of PTPN22 function led to enhanced signaling in T cells expressing a lower avidity self-reactive TCR. In this setting, loss of PTPN22 mediated enhanced proliferation and Th1 skewing. Importantly, expression of the risk variant in association with a lower avidity TCR also increased proliferation relative to PTPN22 non-risk T cells. Together, these findings suggest that, in primary human T cells, *PTPN22* rs2476601 contributes to autoimmunity risk by permitting increased TCR signaling in mildly self-reactive T cells, thereby potentially expanding the self-reactive T cell pool and skewing this population toward an inflammatory phenotype.

## Introduction

Preventing inappropriate immune responses is critical to the maintenance of peripheral tolerance and the prevention of autoimmunity. The risk of autoimmunity stems from a complex interplay of genetic and environmental factors (1, 2). While the greatest risk factor for most autoimmune disease is HLA haplotype (3–5), an individual’s total genetic risk is determined by a combination of gene variants (6–8), many of which cluster near genes responsible for regulating immune responses (9). A leading example of such an autoimmune risk variant is found in the gene Protein Tyrosine Phosphatase Non-Receptor 22 (*PTPN22*).

While the phosphatase encoded by *PTPN22* can function to modulate a range of immune signaling programs, it has been best studied in lymphocyte antigen receptor regulation (10–12). An arginine to tryptophan substitution (R620W) resulting from a single nucleotide polymorphism (SNP; rs2476601) within *PTPN22,* is broadly associated with a large number of autoimmune diseases (13, 14). *PTPN22* R620W, hereafter referred to as the *PTPN22* risk variant, is one of the strongest, non-HLA, genetic risk factors for autoimmune diseases, nearly doubling a carrier’s risk of type 1 diabetes, systemic lupus erythematosus, or rheumatoid arthritis (15–17). As PTPN22 plays a role in multiple autoimmune diseases, it has been argued that PTPN22 potentially highlights a common pathway that alters immune tolerance and might serve as a potential point of intervention to prevent and/or delay disease (18).

In T cells, PTPN22 functions to dephosphorylate key activating tyrosine residues within the proximal tyrosine kinase effectors, LCK and ZAP70, in the TCR signaling pathway (11, 15). This function is mediated in concert with activity through PTPN22’s binding partner, C-terminal Src Kinase (CSK), which also regulates LCK activity (19). Co-association of PTPN22 and CSK results in residence of both proteins within the cytosolic plasma membrane of resting T cells through the association of CSK with another TCR inhibitory protein, PAG (20, 21). The R620W substitution in PTPN22 is located within a proline rich, C-terminal domain and reduces the interaction of PTPN22 with CSK leading to altered localization of PTPN22 (15). The functional outcome of an alteration in PTPN22 protein interactions has led to significant debate as to the mechanism by which the *PTPN22* risk variant contributes to altered T cell function and self-tolerance. Primary T cells isolated from carriers of the *PTPN22* risk variant display significantly weaker responses to TCR engagement, as measured by calcium flux, cytokine release and other metrics (22–25). However, in murine models, expression of the risk variant leads to enhanced TCR responses, as measured by the same metrics in addition to increased T cell proliferation and survival (26–28). In contrast, genetic ablation of *PTPN22* in both mouse or human primary T cells results in enhanced TCR signaling relative to *PTPN22* wild type cells; findings similar to murine risk model yet opposite of the phenotypes described in human carriers of the risk variant (29, 30). These discrepancies in the signaling of primary T cells has led to a continued debate as to whether the human *PTPN22* risk variant mediates a gain- vs a loss-of- function impact on TCR regulation.

Both human carriers and murine models of the *PTPN22* risk variant share important traits. For instance, the risk variant is associated with an increase in memory lymphocyte populations and expanded pro-inflammatory/Th1 T cells and both exhibit increased risk of autoimmune manifestations (25, 27, 31). Thus, it remains surprising that murine and human carriers exhibit these key similarities despite the published differences in the TCR signaling responses described human vs. mouse T cells. Notably, the murine and human T cell populations used for previous studies exhibit fundamental differences. First, most data for human *PTPN22* risk variant phenotype have utilized mature T cells isolated from adult donors, while T cells from murine studies utilize donor mice with minimal environmental exposures. Studies of PTPN22 using human cord blood derived T cells have been limited (18). Also, murine models of the *PTPN22* risk variant are conducted in genetically homogenous, autoimmune-prone strains or models with a fixed TCR repertoire (27, 28, 32, 33), conditions difficult to mimic in a primary human T cell setting. Further, in vitro human T cell line models of the *PTPN22* risk variant lead to discrepant results, indicating that the impact(s) of the variant are context dependent, further complicating efforts to study the variant in human T cells (23, 34).

Recently, Crispr/Cas9 gene editing has allowed for precise manipulation of genetic loci in primary human T cells. Delivery of recombinant ribonucleoproteins (RNPs; (30, 35, 36) containing Cas9 complexed with sequence specific guide RNA sequences) via electroporation into primary T cells efficiently mediates site-specific DNA double stranded breaks (DSB) near a protospacer adjacent motif (PAM) site (37). Such DNA breaks trigger a series of events driving 5’ to 3’ exonuclease activity as the broken DNA ends prepare to modify the break through alternative repair pathways (38, 39). When DSBs are generated in the presence of a single stranded oligo deoxynucleotides (ssODN) (co-delivered with RNP), DNA repair pathways can utilize regions of homology within an ssODN as a repair template for precise genetic modification by homology directed repair (HDR) (40, 41). This approach has previously been used to integrate a premature stop codon in PTPN22 of primary human T cells with high efficiency (30).

In this study, we utilized a combination of HDR-based gene editing and TCR co-delivery in naïve cord blood T cells to establish a platform to better understand the role of the PTPN22 risk variant in primary human T cells. Through Crispr/Cas9 gene editing we find the *PTPN22* risk variant T cells mimicked *PTPN22* KO T cells, with enhanced expression of surface activation markers in response to TCR stimulation. Further, using transgenic TCR stimulation in concert with gene edited cells, we found that the *PTPN22* risk variant accentuates the response of low-avidity primary human T cells to antigen, implying a key role for these events in the progressive loss of T cell tolerance in human autoimmune diseases associated with this common risk allele.

## Materials and Methods

### Primary T cell isolation and culturing

PBMCs were collected from whole (peripheral or cord) blood of consenting donors and cryopreserved. Upon thaw, naive CD4^+^ T cells were isolated by negative selection (Naïve CD4 T Cell Isolation Kit II, Miltenyi Biotech) and cultured in Roswell Park Memorial Institute (RPMI) 1640 media supplemented with 20% FBS, 1x Glutamax (Gibco), and 1mM HEPES (Gibco). Unless otherwise noted, cells were cultured in 5ng/ml recombinant IL-2 (Peprotech). After thaw cells were counted and cultured at 1 million/ml in flat bottom culture plates.

### CRISPR/Cas9 and ssODN reagents

CRISPR gRNAs targeting *PTPN22* exon 14 were commercially synthesized by Synthego (CRISPRevolution sgRNA). ssODNs were commercially synthesized by Integrated DNA Technologies (IDT; Ultramer DNA Oligonucleotides) with phosphorothioate linkages between the first and final 3 base pair sequences. gRNA complexes were mixed with Cas9 nuclease (IDT) at a 1.2:1 ratio and delivered with or without ssODNs into cells by electroporation. *PTPN22* targeting 20bp gRNA sequence used for all editing: AATGATTCAGGTGTCCGTAC.

### Antigen specific TCR Lentiviral Vectors (LV-TCRs)

Antigen specific TCRs utilized in this study have been previously described ((42, 43), T1D2 herein described as “Lower Avidity TCR” (L-TCR) with an EC50 of 0.97 μg/mL and T1D5-2 herein described as “Higher Avidity TCR” (H-TCR), EC50 0.09 μg/mL. TCR coding sequences were cloned into lentiviral backbones using InFusion HD (Takara), with oligonucleotides synthesized by IDT, and a cis-linked (T2A) eGFP was inserted into the 3’ end of the alpha constant region. LV-TCR plasmids were based on the pRRL backbone derived from pRRLSIN.cPPT.PGK-GFP.WPRE (plasmid #12252, Addgene). The DNA sequences of all plasmids were verified by Sanger sequencing performed by GENEWIZ. Methods for LV production have been previously described (44). Lentivirus stocks were tittered by rtPCR and found to produce similar rates of transduction when administered at the same MOI.

### Gene Editing

After thaw, cells were activated with CD3/CD28 Activator Beads (Gibco). After 2 days beads were magnetically removed and cells re-plated without changing media or adjusting cell number. One day later, cells were electroporated with editing reagents. Prior to electroporation, cells were washed with PBS and resuspended in Lonza P3 buffer. 5μg of complexed RNP and 100 pmol ssODN per 1×10^6^ cells was added to the resuspension so that the final cell density was 5 × 10^7^ cells/ml. Cells were electroporated with a Lonza 4-D Nucleofector, using program DN-102 with 16-well 20µL Nucleocuvette strips or 100µL Nucleocuvette vessels, and then transferred into pre-warmed cell culture medium with IL-2 (unless otherwise noted). For samples transduced with lentivirus, virus was added to the culture 1 day after bead stim addition at an MOI of 5. After editing, cells were maintained in media identical to pre-editing conditions (unless otherwise noted). Cells were counted at least every two-days using Count Bright absolute counting beads (Thermo Fisher Scientific) and split to maintain cell density of 1 to 2 million/ml. Following expansion cells were counted, washed 2 times with PBS, and rested at 1.5 million/ml for 24 hours in cytokine free media consisting of RPMI 1640 media with 10% FBS, 1x Glutamax (Thermo Fisher Scientific), and 1mM HEPES. Cells were re-counted prior to stimulation. For some experiments, cells were frozen after 24- hour cytokine free rest, then thawed/ counted prior to stimulation.

### ddPCR

Quantification of HDR and NHEJ rates in edited human CD4^+^ T cells was obtained using a droplet digital PCR, dual-probe competition assay. All probes were ordered from Sigma Aldrich with a 3’ Black Hole 1 Quencher. Probes specific to sequences generated by HDR insertion of SNP edits or stop codons were labeled with a 5’ FAM reporter and used in tandem with a 5’ HEX labeled probe specific to wild type (WT) sequences. Editing was measured after generating droplets with 50ng of genomic DNA (gDNA), both HDR-FAM and WT-HEX probes, and primers to the editing locus producing amplicons of <500bp (1× assay, 900-nM primers and 250-nM probe) using ddPCR supermix for probes (no deoxyuridine triphosphate [dUTP]) (Bio-Rad). Reference reactions were simultaneously performed using a 5’ HEX labeled probe/ primer combination targeting the houskeeping gene B2M. Droplets were generated with the QX200 Droplet Generator (Bio-Rad) and amplified. All samples were run in triplicate and averaged. Fluorescence was measured using the QX200 Droplet reader (Bio-Rad) and analyzed using Quantasoft software. Editing rates were calculated as the relative frequency (%) of FAM+ corresponding to %HDR, HEX+ corresponding to %No Event, and reference – (FAM+HEX) corresponding to %NHEJ.

For ddPCR analysis of cocultured PTPN22 risk/ non-risk edited cells HDR (risk) -FAM and HDR (non-risk) -HEX probes with locked nucleic acid (LNA) modifications were utilized. LNA modified nucleotides were integrated at the SNP nucleotide and the adjacent 5’ and 3’ nucleotides to increased probe discrimination. Risk/ non-risk probe reads were compared to a B2M reference reaction as above.

### Western blotting

All western blots were performed on lysates from human umbilical cord blood derived CD4 T cells that had been either mock or gene edited, expanded 7 days in cytokine supplemented media, and subjected to 24 hours rest in cytokine free media. Cells were lysed in 1x RIPA lysis buffer on ice for 10 minutes then clarified by centrifugation. Concentration of clarified lysate was determined by BCA assay (Pierce), diluted, and suspended in 1x LDS Sample Buffer (Invitrogen). 10μg of lysate was run on 4-12% Bis-Tris NuPAGE gels in 1x MOPS buffer (Invitrogen). Protein was transferred to nitrocellulose in 1x Transfer Buffer (Invitrogen) and 10% methanol. Non-specific binding was minimized with a 1-hour RT incubation in Odessey LI-COR Blocking Buffer. Primary antibodies were stained at 1:1000. PTPN22 primary stain was for at least 12 hours at 4°C and Actin was stained at RT for 40 minutes. Primary antibodies used were from Cell Signaling Technology: PTPN22 (D6D1H) and Actin (8H10D10). After primary stain membranes were washed with 1x TBST and incubated with secondary antibodies at 1:10,000 for 30 minutes at RT. Stained blots were washed and imaged on an Odyssey Infrared Imaging System (LI-COR Biotech.). Western blot quantifications were performed with ImageJ software.

### Plate bound anti-CD3 stimulation and Th1 skewing

Stimulation plates were made in 96-well flat bottom culture plate. 100ul of PBS supplemented with LEAF purified anti-CD3 (OKT3, Biolegend) at 0.25ug/ml was added to each well and incubated at least 12 hours at 4°C. The plate was then emptied, and wells were given 100ul of cytokine free T cell media. After cells were edited, expanded for 7 days, and rested 24 hours in cytokine free media, 100ul of cells at 2 million/ml were added to each well. Plates were incubated at 37°C for up to 48 hours.

For Th1 skewing of edited T cells, cells that had been edited and rested were transferred to 96-well flat bottom culture plates, coated with anti-CD3 as discussed above. These cells were cultured for 3 days in media containing 20 ng/ml IL-12 (Peprotech), 50 ng/ml IL-2, 40 μg/ml anti-IL-4 (MP4-25D2, Fisher Sci), and 500 ng/ml soluble CD28 (28.2, BioXcell) to skew them to a Th1 phenotype. Non-skewed controls were cultured in media that contained IL-2 and CD28.2 only.

### Peptide stimulation of antigen specific edited T cells

LV-TCRs used in this study are HLA-DR0401 restricted and recognize the 20-mer peptide sequence: QLYHFLQIPTHEEHLFYVLS (IGRP- p39 position 305-324). For stimulation of cells expressing LV-TCRs, antigen presenting cells (APCs) were PBMCs from an allogeneic HLA-DR0401 donor irradiated with 5,000 rads. APCs were then mixed with peptide diluted in complete media (final peptide concentration 50ng/ml) for 2 hours. Co-cultures were incubated in 96 well u-bottom plates with 200K APCs and 50K T cells per well at 37°C for up to 3 days in complete media containing no exogenous cytokines. APCs for all experiments were from the same PBMC donor.

### Flow cytometry and gating strategies

Flow cytometric analysis was performed on an LSR II flow cytometer (BD Biosciences) and data was analyzed using FlowJo software (Tree Star). Cells were stained with LIVE/DEAD Fixable Near-IR Dead Cell Stain Kit (Fisher Sci), as per the manufacturer’s instructions and cells were stained with fluorescence labeled antibodies for 30 minutes on ice. Antibodies used in this study include from Biolegend: CD3 (SK7), CD4 (RPA-T4), CD69 (FN50), CD25 (M-A251), CD71 (CY1G4), IFNγ (4S.B3), IL-2 (MQ1-17H12), and anti-mouse TCR-β (H57-597); and from Cell Signaling: phospho-S6 kinase (D57.2.2E).

All surface antibodies were used at a dilution of 1:100, antibodies against cytokines were used at a dilution of 1:200, and those staining phospho-sites were used at a dilution of 1:50. Gating order proceeded: lymphocytes -> singlets -> live cells. Surface stains of other markers were subsequently gated on CD4^+^/TCR^+^ cells, then the marker of interest. For pS6 staining, cells were fixed with a final concentration of 2% PFA for 12 minutes at 37°C. Cells were then washed and permeabilized with BD Perm Buffer III for at least 30 minutes at -20°C. Cells were then washed wand stained as described above. For cytokine staining, after 3 days of stimulation cells were stimulated with 50ng/ml PMA, 1ug/ml Ionomycin, and 1ug/ml monensin for 5 hours, then fixed/ permed with BD Cytofix/Cytoperm prior to staining for cytokine production. For assessment of proliferation, peptide stimulated antigen specific T cells were labeled with CellTrace Violet proliferation dye (Fisher Sci.) prior to co-culturing with APCs; number of antigen specific T cells in each edited sample was determined daily by addition of Countbright Absolute Counting Beads (Fisher Sci.) in flow cytometry samples.

### Statistics

Statistical analyses were performed using GraphPad Prism 9 (GraphPad). For all testing of gene edited cells, due to the low variability in culturing conditions and lack of obvious skewing, data was assumed to maintain a normal distribution. P-values in multiple comparisons were calculated using one-way ANOVA with the Tukey correction; p-values in comparisons between two groups were calculated using a paired two-tailed t test. Values from combined independent experiments are shown as mean ± SEM.

### Study Approval (Human Subjects)

For gene editing experiments using adult PBMCs, human donor leukopaks were purchased from the Fred Hutchinson Cancer Research Center, which were obtained from consenting donors under an IRB-approved protocol and cryopreserved. For gene editing experiments using umbilical cord blood derived PBMCs, cord units were purchased from the Bloodworks Northwest, which were obtained with consent under an IRB-approved protocol and cryopreserved. After collection, all samples were de-identified for the protection of human blood donors.

## Results

### Generation of PTPN22 risk and non-risk edited T cells with Crispr/Cas9 gene editing

In this study, we sought to identify methods to utilize HDR-based gene editing to study alterative *PTPN22* alleles within identical populations of primary human T cells. Notably, previous work has shown that altering expression levels of PTPN22 and/or its regulatory proteins can produce potentially contradictory results (23, 34). Additionally, *PTPN22* contains several splice variants that might modulate the impact of the risk variant (45). Therefore, we sought to utilize an editing platform that would permit us to introduce either a control, risk, or knock out sequence via direct modulation of the endogenous SNP locus, without the addition of exogenous promoters or selection agents that may alter natural PTPN22 expression. To accomplish this goal, we expanded on previously reported methods to generate PTPN22 gene disrupted primary human T cells (30) to established an HDR-based editing platform. We combined delivery of Crispr/Cas9 with ssODN HDR templates designed to alter the coding region of Exon 14 of *PTPN22.* Use of Crispr/Cas9 RNPs with short ssODN repair templates to alter the DNA of primary cells has been shown to be highly efficient, provided the nuclease cuts near the desired edit site (46). A naturally existing Crispr/Cas9 gRNA PAM site is positioned in Exon 14 of *PTPN22* to allow an RNP induced DSB immediately 5’ of the SNP nucleotide (Fig. 1, A). Also, previous work has shown that homology arm design of ssODN repair templates may impact editing rates when used with a Crispr/Cas9 RNP (40). We tested several ssODN repair template designs with our *PTPN22* RNP for optimized introduction of a 2bp coding alteration to generate the PTPN22 risk variant (Fig. 1, A; Fig. S1, A).

**Figure 1.**
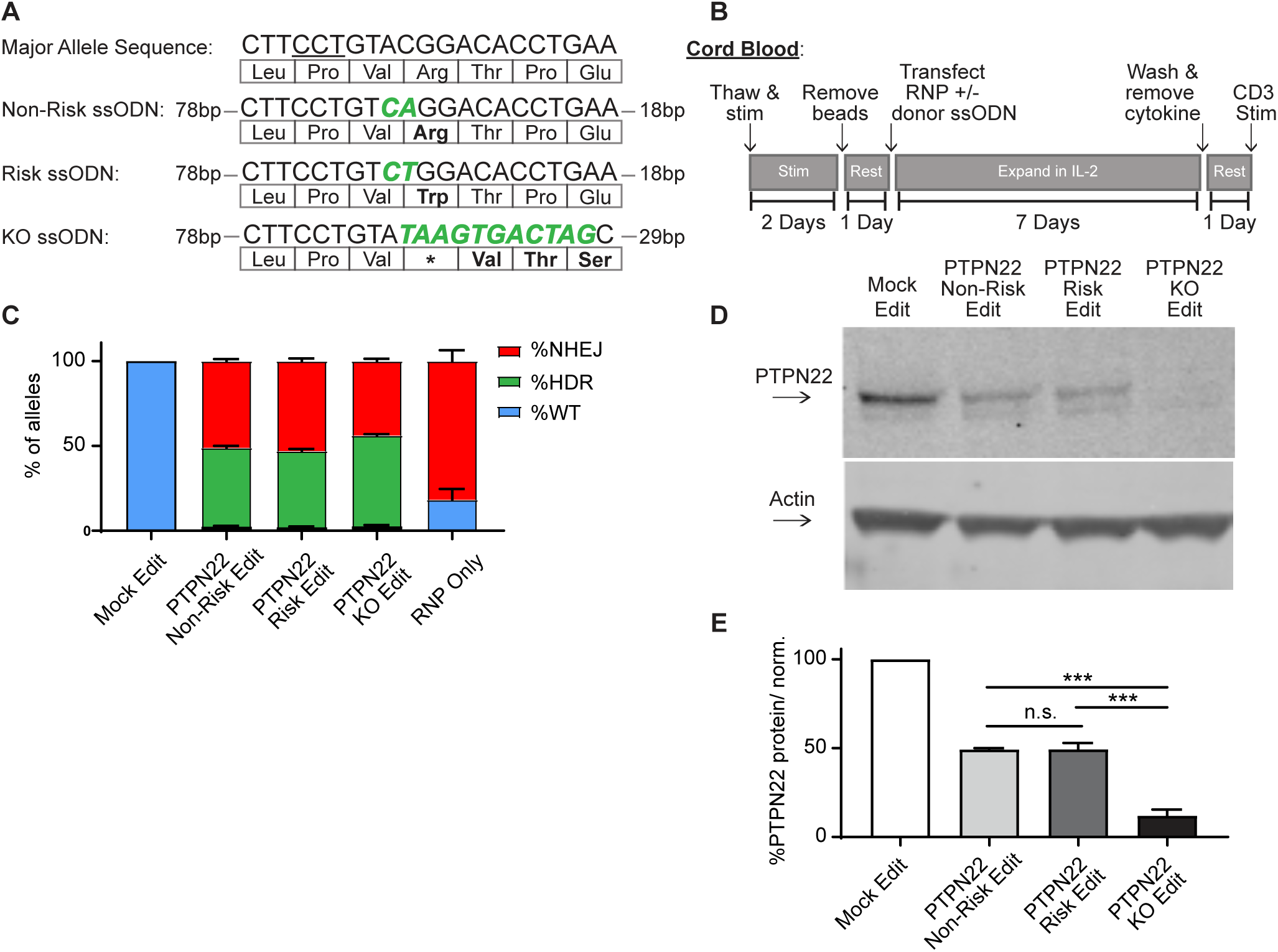
Gene editing PTPN22 SNP variants in cord blood CD4 T cells produces comparable populations. CD4 T cells were isolated from human umbilical cord blood for Crispr/Cas9 gene editing. (A) ssODN repair templates electroporated with *PTPN22* targeting Crispr/Cas9 RNP resulting in differently coded alleles upon HDR. Underlined nucleotides in major allele sequence indicate RNP PAM site. PTPN22 coding alterations for each population denoted with bolded/ colored nucleotides, resulting changes to amino acid sequence displayed below. (B) Workflow used to produce and assay *PTPN22* edited CD4 T cells and corresponding controls. (C) ddPCR analysis of editing outcomes in *PTPN22* edited cells using corresponding ssODN repair templates or RNP alone. n=4 independent human donors. (D) Representative western blot of PTPN22 expression in mock edited, PTPN22 non-risk edited, PTPN22 risk edited, and PTPN22 knockout edited cord blood CD4 T cells from the same human donor. (E) Quantified PTPN22 protein expression relative to Actin and normalized to unedited values from the same T cell donor. n=3 independent human donors, matched one-way ANOVA with Tukey’s correction. For summary graphs lines and error bars represent mean +/- SEM. RNP - ribonucleoprotein, ODN or ssODN - single stranded oligo-deoxynucleotide, NHEJ - non-homologous end joining, HDR - homology directed repair. All data is from at least 2 independent experiments. *** p<0.001.

Importantly, this 2bp edit blocks subsequent binding of the RNP, preventing cleavage of the repaired locus. We found that in adult peripheral blood CD4 T cells, a 120bp ssODN design with a long 5’ homology arm (ssODN V3, 5’ arm 89bp/ 3’ arm 29bp) efficiently introduced the desired coding sequence alterations (Fig. S1, B), in line with previous findings (40). Using this design, we created three ssODN repair templates that worked efficiently in association with the PTPN22 RNP to produce 3 gene editing outcomes: non-risk edited *PTPN22*, risk edited *PTPN22*, and a PTPN22 KO ssODN that seamlessly introduces a stop codon into all 3 reading frames of the RNP cut site (Fig. 1A).

Natural carriers of the *PTPN22* risk variant have altered peripheral T cell responses, and mouse models indicate that variant expression impacts thymic T cell selection as well as peripheral T cell activation and function (27). Thus, human T cells isolated from subjects with the risk variant are likely to be previously impacted by diverse developmental, age-related and environmental exposures (18). Therefore, we reasoned that using naïve, cord blood derived CD4 T cells isolated from non-risk human donors, which are minimally impacted by previous environmental events, would provide the most relevant cell population for our study and generate the most informative dataset with respect to impact(s) of the variant on human T cell function. Using our *PTPN22* RNP and ssODN repair templates with a gene editing workflow (Fig. 1B), we edited human cord blood derived naïve CD4 T cells from multiple *PTPN22* non-risk donors to concurrently model the *PTPN22* risk variant, control edited, and knock-out edited cells. Comparing edited populations to mock edited cells (which received Cas9 without gRNA and PTPN22 KO ssODN), all gene edited populations showed similar viability, regardless of editing reagents used, with slight reduction in viability observed all T cell populations receiving electroporation (Fig. S1, C). Digital droplet PCR (ddPCR) analysis demonstrated nearly identical gene editing rates in cells edited with the *PTPN22* non-risk vs. risk ssODN (Fig. 1, C). Both edited populations exhibited average HDR rates of ∼50%, and no detectable editing was present in mock edited populations. HDR rates in PTPN22 KO edited cells were slightly higher, likely due to a lack of nucleotide mismatches between the ssODN template and the digested *PTPN22* locus. As expected, beyond HDR events, NHEJ events accounted for most of the remaining editing outcomes in cells modified in the presence of ssODNs. NHEJ events were absent or dominant, in mock and RNP-only edited populations, respectively. Western blot analysis confirmed that, *PTPN22* non-risk and risk edited populations retained equivalent expression levels of PTPN22 (with a reduction in total PTPN22 protein compared with mock edited cells reflecting the NHEJ mediated loss of protein expression in the absence of successful HDR). As expected, PTPN22 KO populations exhibited near total absence of protein expression (Fig. 1D-E).

Wild type PTPN22 alleles were nearly undetectable in edited populations (<1% of total alleles), indicating that resulting genotypes of edited T cells can be assumed to a mixture of biallelic HDR, biallelic NHEJ (PTPN22 KO), or HDR and NHEJ on opposite alleles (hemizygous). RNP based editing can induce large deletions that may bias sequencing of editing outcomes (47), making *accurate* single cell profiling challenging. As HDR rates in our approach were identified using probe based ddPCR detection, our approach is resistant to this type of bias. With ∼50% HDR rates and ∼50% NHEJ rates, the edited populations would at a minimum be half biallelically edited and half full PTPN22 KO. Of note, previous single cell studies in primary cell types (48, 49) including primary T cells (49), demonstrate that monoallelic HDR is far more frequent than biallelic HDR. Therefore, it is reasonable to assume the predominant editing outcome in our studies is hemizygosity with an undetermined frequency of biallelic editing events. However, as an independently edited control population is also utilized in our analysis, the effects of PTPN22 KO alleles (monoallelic and biallelic) are accounted for when PTPN22 risk edited cells are compared to non-risk edited populations. Thus our editing approach assures that our findings in PTPN22 risk edited populations can be attributed to the expression of differently coded PTPN22 alleles, not PTPN22 loss of function.

### PTPN22 risk edited T cells mimic PTPN22 KO cells

Following generation of T cells expressing equivalent levels of the *PTPN22* risk, non-risk, or PTPN22 KO T cells from the same donor, we tested how alterations in *PTPN22* coding impacted responses to TCR stimulation. After gene editing and expansion, T cells were stimulated with plate bound anti-CD3 for 6 to 48 hours and sequentially evaluated for expression levels of key surface activation markers. As all *PTPN22* edits produced some loss of PTPN22 expression, and PTPN22 is responsible for restraining TCR activation, it was unsurprising to find that mock edited cells had significantly less expression of the activation markers CD69, CD25, and CD71 compared with gene edited populations (Fig. 2, A-C). However, it was striking that, while PTPN22 KO populations expressed the highest levels of these activation markers, the *PTPN22* R620W risk-edited cells, in comparison with non-risk edited cells, exhibited increased expression of CD25 and CD71 at 48 hours post stimulation and a trend toward increased CD69 expression as early as 6 hours post stimulation (Fig. 2, A-C). Together, this data strongly suggest that TCR signaling strength is increased in cells expressing the PTPN22 risk variant, and, that this phenotype is most similar to PTPN22 KO T cells.

**Figure 2.**
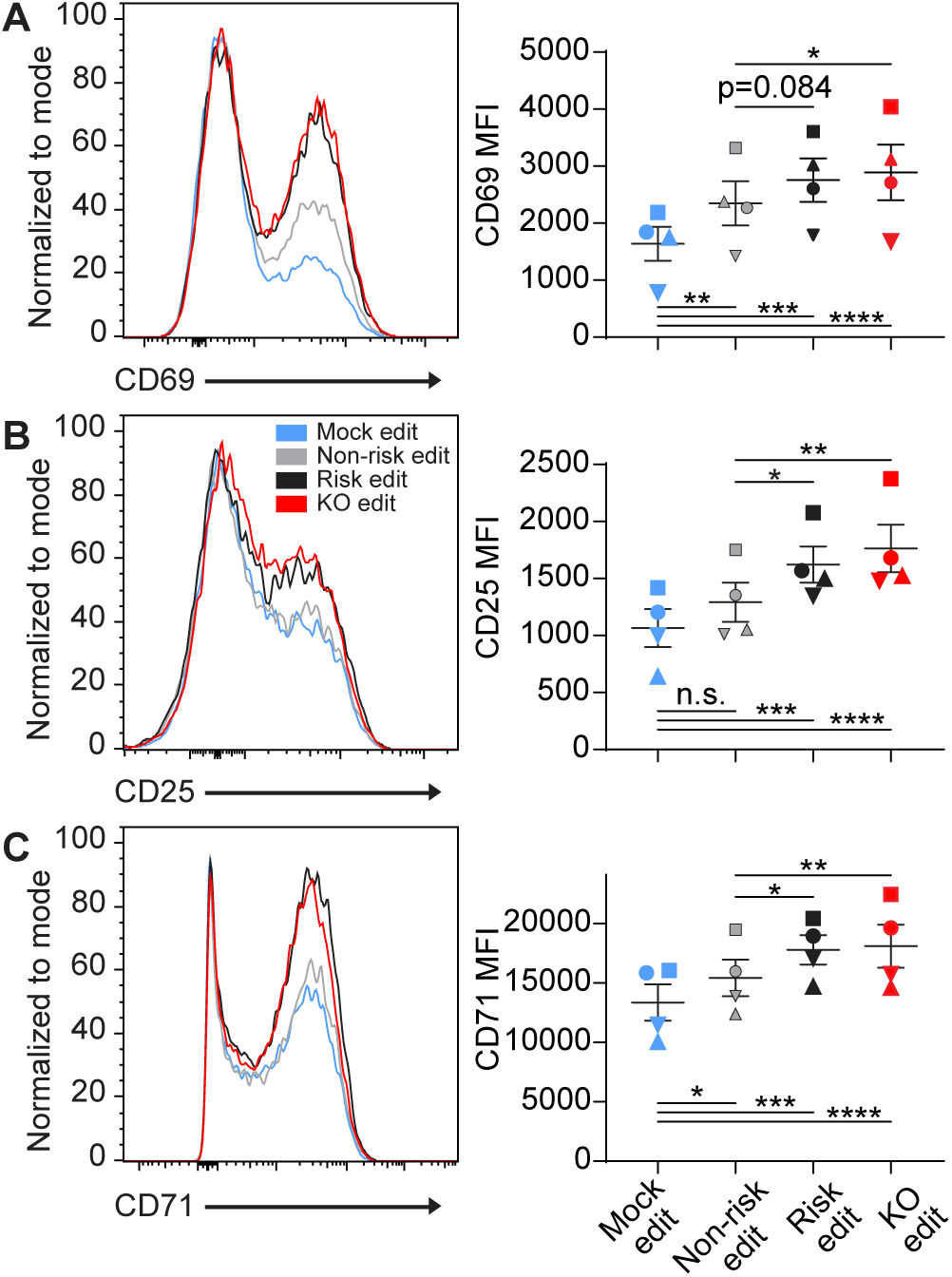
PTPN22 risk edited T cells mimic PTPN22 KO in response to TCR stimulation. Cord blood CD4 T cells from human donors were gene edited as described in Figure 1, then stimulated with plate bound anti-CD3 for up to 48 hours. (A-C) Representative flow overlays for CD69 (A), CD25 (B), and CD71 (C) with summary data for different editing conditions from the same donor. CD69 summary data reflects mean fluorescent intensity at 6 hours post CD3 stimulation, while CD25 and CD71 represent median values at 48 hours post stimulation. For all summary data n=4 independent donors (shapes correspond to individual donors), matched one-way ANOVA with Tukey’s correction. Lines and error bars represent mean +/- SEM. All data is from 2 independent experiments. * p<0.05, ** p<0.01, *** p<0.001, **** p<0.0001.

The *PTPN22* risk variant has been reported to modulate CD4 T cell populations, with some studies suggesting increased proportions of Th1 (25, 31, 32) populations in risk carriers. To examine if an altered PTPN22 status is sufficient to modulate CD4 T cell skewing outcomes, *PTPN22* edited cord blood T cells were cultured with plate-bound anti-CD3, soluble CD28 and IL-12 to produce Th1 CD4 T cells. Three days after stimulation we found most cells producing Th1 associated cytokines, however no differences in the number of cells producing IFNγ or IL-2 were observed based upon edits made to *PTPN22* (Fig. S2, A-B). This lack of difference in Th1 skewing is consistent with findings in a similar study using the murine *PTPN22* risk variant model (31).

### PTPN22 regulates signaling from low avidity TCRs in human T cells

PTPN22 is a key regulator of TCR signaling and in murine models it plays a critical role in repressing T cell activation mediated by weak TCR agonists, but is dispensable in regulating responses to strong TCR agonists (32). These findings suggested that perturbations in PTPN22 activity might impact autoimmune risk via enriching T cells expressing low-avidity self-reactive TCRs that escaped deletion during thymic development and remain capable of recognizing self-antigens. Thus, as our gene editing model indicated that the *PTPN22* risk variant produces a phenotype similar to a PTPN22 knockout, we next examined whether PTPN22 knockout modified the response of human T cells expressing TCRs with differing avidity for an identical self-antigen.

Previous work has detailed the sequencing of multiple TCRs from T1D patients that are reactive against the pancreatic islet autoantigen IGRP (42, 43), and recent work has shown that gene edited primary human T cells can be rendered antigen specific through the lentiviral delivery of these TCRs prior to electroporation with RNPs (50). To study the impact of PTPN22 loss of function on antigen specific TCR responses, we transduced human cord blood CD4 T cells with lentiviral vectors (LV) encoding for alternative TCRs that respond to the same 20mer epitope of IGRP when presented by HLA-DRB1 04:01; and subsequently modified these LV transduced populations using the HDR-based gene editing (Fig. 3, A). Addition of LV transduction prior to gene editing did not impact gene editing outcomes or rates of HDR (Fig. S3, A). Transgenic TCR chains were engineered to utilize the murine constant regions allowing staining with murine TCR specific antibodies (42) and limiting potential cross pairing with endogenous TCR chains. To track LV transduced cells, constructs were modified to express cis-linked GFP simultaneously with TCR expression. LV transduction rates were found to be largely identical regardless of TCR sequence utilized and were titrated to transduce approximately 25% of the culture, helping ensure ∼1 viral copy per transduced cell to achieve similar levels of TCR expression in all transduced cells. After gene editing, expansion, and rest, gene edited/ TCR+ T cells were stimulated via addition of irradiated PBMCs from an allogeneic HLA-DRB1 04:01 donor that had been pre-incubated with cognate IGRP peptide (Fig. 3, B). By labeling TCR+ cord blood T cells with a proliferation dye, then stimulating with peptide loaded APCs, we demonstrated that these self-reactive TCRs responded to the IGRP peptide with markedly different rates of cell proliferation over a 3-day stimulation (Fig. 3, C). The two TCRs, which we will term: “Lower Avidity TCR” or L-TCR, and “Higher Avidity TCR” or H-TCR, were found to trigger significantly different rates of proliferation across a range of peptide doses, consistent with their reported avidity (Fig. 3, C)(43). As the highest peptide dose reliably produced efficient proliferation in all donors tested, we used this dose for activation of *PTPN22* edited cells.

**Figure 3.**
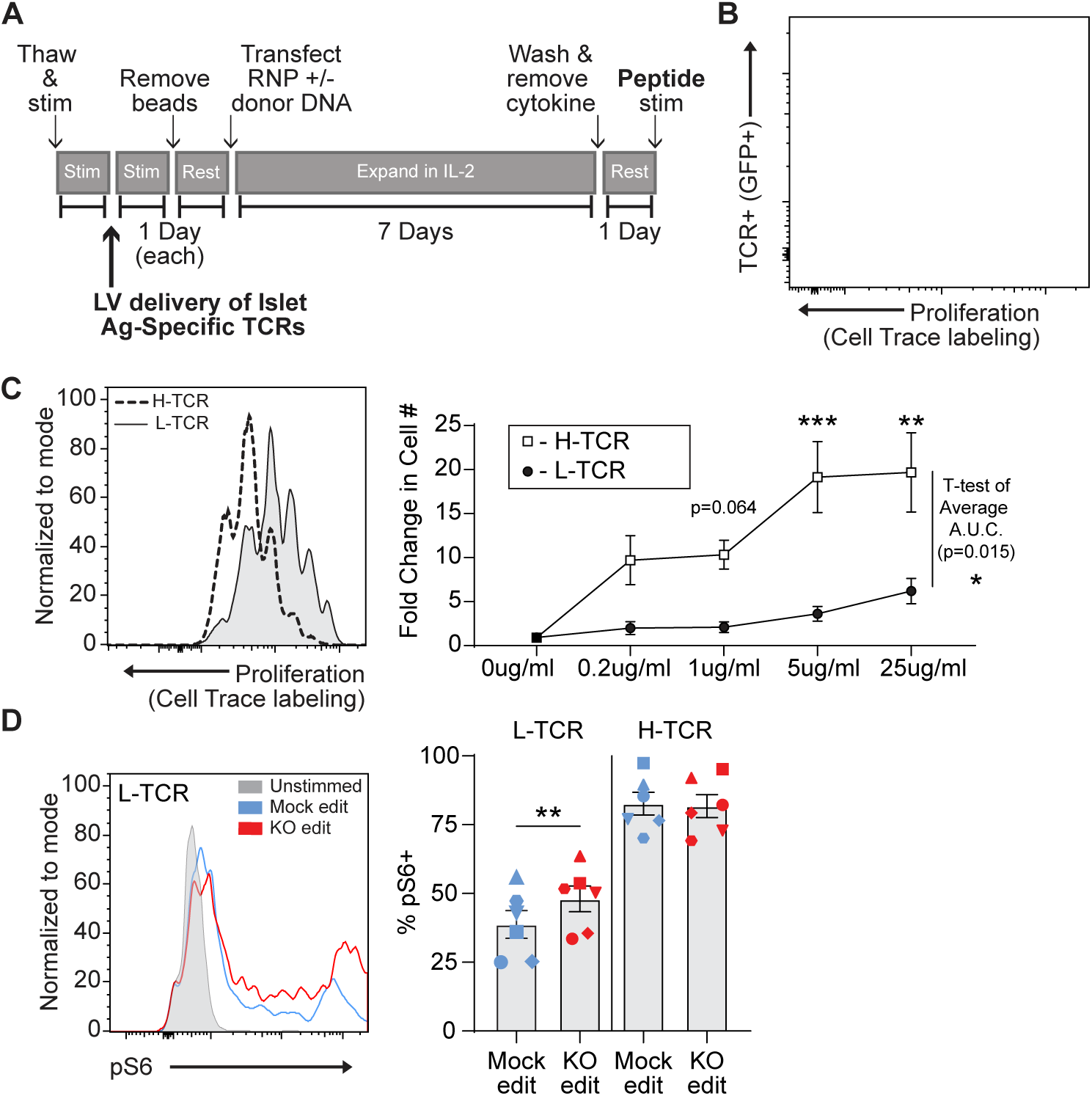
Antigen specific PTPN22 KO T cells have increased phosphorylated S6 in response to low avidity TCR engagement low avidity TCR stimulation. Cord blood CD4 T cells from human donors were transduced with lentivirus coding for antigen specific TCRs prior to gene editing as described in Figure 1, then stimulated with cognate peptide. (A) Workflow shown in figure 1, modified to allow both gene editing of the *PTPN22* locus and lentiviral delivery of antigen specific TCRs. (B) Representative flow plot of H-TCR+ T cell proliferation after 3 days of peptide stimulation. (C) Representative flow overlay of 3-day peptide induced proliferation caused by either H- TCR stimulation or L-TCR stimulation in T cells from the same donor and summary data of 3-day T cell proliferation (fold change over input cell number) at various peptide doses in cells expressing transgenic TCRs. Proliferation induced by TCRs was compared by computing area under the curve (A.U.C.) for each donor at all peptide dosed tested, then performing a paired T test. Proliferative effect of each TCR at individual peptide concentrations was analyzed by a paired 2-way ANOVA with a Sidak multiple comparison test. (D) Representative flow overlay of different editing conditions from the same donor expressing L-TCR and summary data of flow values for phosphorylated S6 kinase in edited cells expressing transgenic TCRs, n=6 independent donors (shapes correspond to individual donors), paired T test by TCR expressed. Columns and error bars represent mean +/- SEM. All data is from 3 independent experiments. * p<0.05, ** p<0.01, *** p<0.001.

To first examine how loss of PTPN22 impacted antigen-specific TCR signaling, we generated mock-edited vs. PTPN22 KO cord blood CD4 T cells expressing either the L-TCR or H-TCR and assessed TCR-mediated, downstream phosphorylation events via flow cytometry. In cells expressing H-TCR, PTPN22 KO had no impact on the percent of pS6+ cells. In contrast, for cells expressing L-TCR, the loss of PTPN22 resulted in significantly greater proportion of pS6+ cells (Fig. 3D).

We next determined if the early signaling differences found in PTPN22 KO cells expressing L-TCR (vs H-TCRs) led to differences in activation phenotype. Two days post-peptide stimulation, L-TCR expressing PTPN22 KO T cells, in comparison to mock edited T cells from the same cord blood donor also expressing L-TCR, expressed higher levels of activation markers including CD69, CD25, and CD71 (Fig. 4, A-C). In contrast, H-TCR expressing PTPN22 KO T cells exhibited no differences in comparison to control T cells. Together, these results demonstrate that loss of PTPN22 function preferentially permits enhanced signaling in response to low avidity TCR engagement in primary human T cells.

**Figure 4.**
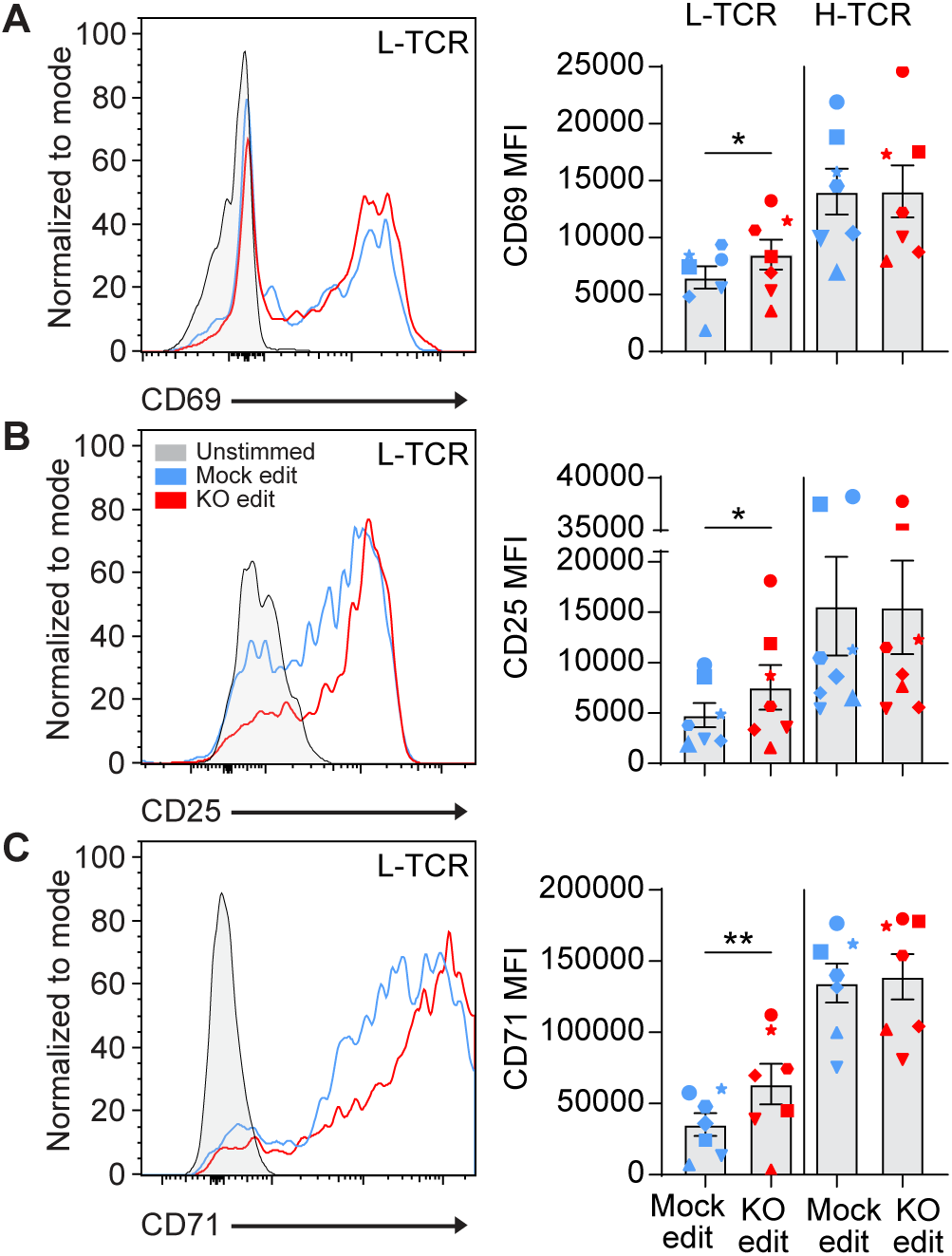
L-TCR stimulation of PTPN22 KO edited T cells produces enhanced surface activation marker expression. Cord blood CD4 T cells from human donors were edited as described in Figure 3, then stimulated with peptide loaded APCs for up to 48 hours. (A-C) Representative flow overlays and summary data for CD69 (A), CD25 (B), and CD71 (C) expression in mock edited and PTPN22 KO populations from the same donor expressing the L-TCR. CD69 summary data reflects mean fluorescent intensity at 24 hours post peptide stimulation, while CD25 and CD71 represent median values at 48 hours post stimulation. For all summary data n=7 independent donors (shapes correspond to individual donors), paired T test by TCR expressed. Columns and error bars represent mean +/- SEM. All data is from 4 independent experiments. * p<0.05, ** p<0.01.

### PTPN22 deficiency promotes proliferation and IFNγ secretion by low avidity T cells

We next determined the functional outcomes associated with the increased activation of L-TCR+ PTPN22 deficient T cells and whether potential changes might predict an increased risk for autoimmunity. L-TCR and H-TCR expressing, mock edited or PTPN22 KO cord blood T cells were stimulated for 3 days with cognate peptide. L- TCR+ PTPN22 KO cells had significantly increased proliferation responses relative to mock edited cells expressing L-TCR (Fig. 5A). In contrast, H-TCR+ T cells showed identical rates of proliferation in response to antigen, regardless of the presence or absence of PTPN22 (Fig. 5B).

**Figure 5.**
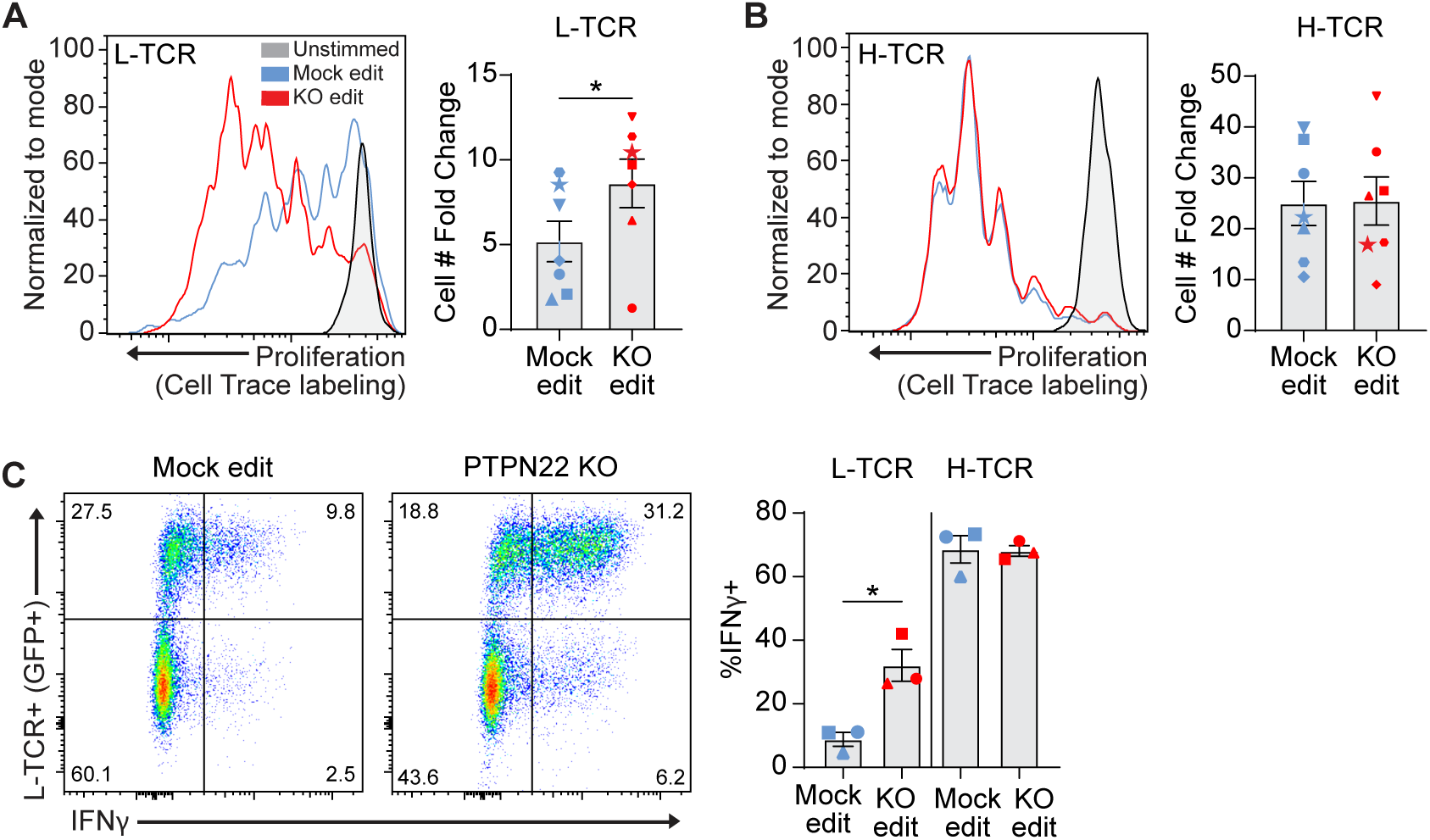
PTPN22 KO T cells show increased proliferation and IFNy secretion in response to low avidity TCR engagement L-TCR stimulation. Cord blood CD4 T cells from human donors were edited as described in Figure 3, then stimulated with peptide loaded APCs for 72 hours. (A) Representative flow overlay of proliferation in L-TCR+/ *PTPN22* gene edited cells from the same donor and summary data of T cell proliferation (fold change over input cell number), n=7 independent donors (shapes correspond to individual donors), from 4 independent experiments. (B) Representative flow overlay of proliferation in H-TCR+/ *PTPN22* gene edited cells from the same donor and summary data of T cell proliferation (fold change over input cell number), n=7 independent donors (shapes correspond to individual donors), from 4 independent experiments. (C) Representative flow plots of IFNy expression in mock edited or PTPN22 KO cells transduced with L-TCR and summary data of %*PTPN22* edited/TCR+ cells that secrete IFNy after 3 days of peptide stimulation in cells expressing either the L-TCR or H-TCR. n=3 independent donors (shapes correspond to individual donors), from 2 independent experiments). All summary data analyzed by paired T test. Columns and error bars represent mean +/- SEM. All data is from 2-4 independent experiments. * p<0.05.

A common feature of autoimmunity is inappropriate generation of proinflammatory cytokines, that drive diverse pathologies in various tissues. While, as previously noted, *PTPN22* status had no impact on the generation of an IL-12 driven Th1 phenotype in association with CD3/CD28 engagement (Fig S2, A-B), we wanted to determine whether *PTPN22* status might impact the generation of a Th1 phenotype in association with TCR stimulation in response to cognate antigen. We found that, three days after co-culture of peptide loaded APCs with PTPN22 KO cells expressing either L- TCR or H-TCR produced significant levels of IFNγ (Fig. 5C). Strikingly, while the proportion of IFNγ secreting H-TCR+ cells was not affected by PTPN22 KO, the proportion of L-TCR+ cells that generated IFNγ were dramatically increased by loss of PTPN22. A similar trend was found with IL-2 secretion (Fig S4, A-B; p=0.068). Together these findings indicate that a key phenotypic outcome of PTPN22 deficiency is expansion of weakly reactive T cells and preferential skewing of these cells toward a proinflammatory phenotype.

Because *PTPN22* risk variant edited T cells exhibited a phenotype similar to PTPN22 KO T cells in response to anti-CD3 stimulation, we next determined whether risk variant edited cells exhibited increased proliferation in response to cognate antigen. Generation of *PTPN22* risk edited cells leads to a reduction in PTPN22 expression due to competing HDR and NHEJ events (Fig. 1). Thus, to control for levels of HDR edited vs. gene disrupted *PTPN22* alleles, we designed our experiments to assess the functional outcome of TCR engagement within a pool of risk vs. non-risk edited cells. To achieve this comparison, cord blood T cells were LV transduced to deliver the L-TCR and subsequently edited (as in Figs. 1A, 3A) to generate *PTPN22* risk edited or non-risk edited populations, each expressing L-TCR (Fig 6A). Next, edited populations were mixed in equal proportions, FACS sorted to isolate TCR+ cells, and labeled with a proliferation dye. This mixed population was then used to test the hypothesis that, in response to stimulation with cognate antigen, *PTPN22* risk edited cells would become the dominant genotype within the proliferating cell population. gDNA based ddPCR analysis demonstrated equal proportions of *PTPN22* risk and non-risk edited alleles (Fig. 6B-C). The mixed population was then stimulated using peptide loaded APCs for 3 days and proliferating (Fig. 6B) vs non-proliferating cells were isolated by FACS sorting (Fig. S5, A-B). ddPCR revealed nearly twice as many *PTPN22* risk edited over non-risk edited alleles within the proliferating cell population (Fig. 6B-C). Together, the data support a hypothesized model (Fig. 6A) wherein human T cells expressing the PTPN22 risk variant exhibit increased proliferation in response to low avidity TCR engagement. Interestingly, gDNA taken from stimulated but non-proliferating TCR+ cells also showed a partial increase in *PTPN22* risk edited alleles relative to input although less robust than among proliferating cells (Fig. S5, A-B). This observation suggests that increased TCR signal strength in risk variant T cells may also promote enhanced cell survival, a finding that will require additional study.

**Figure 6.**
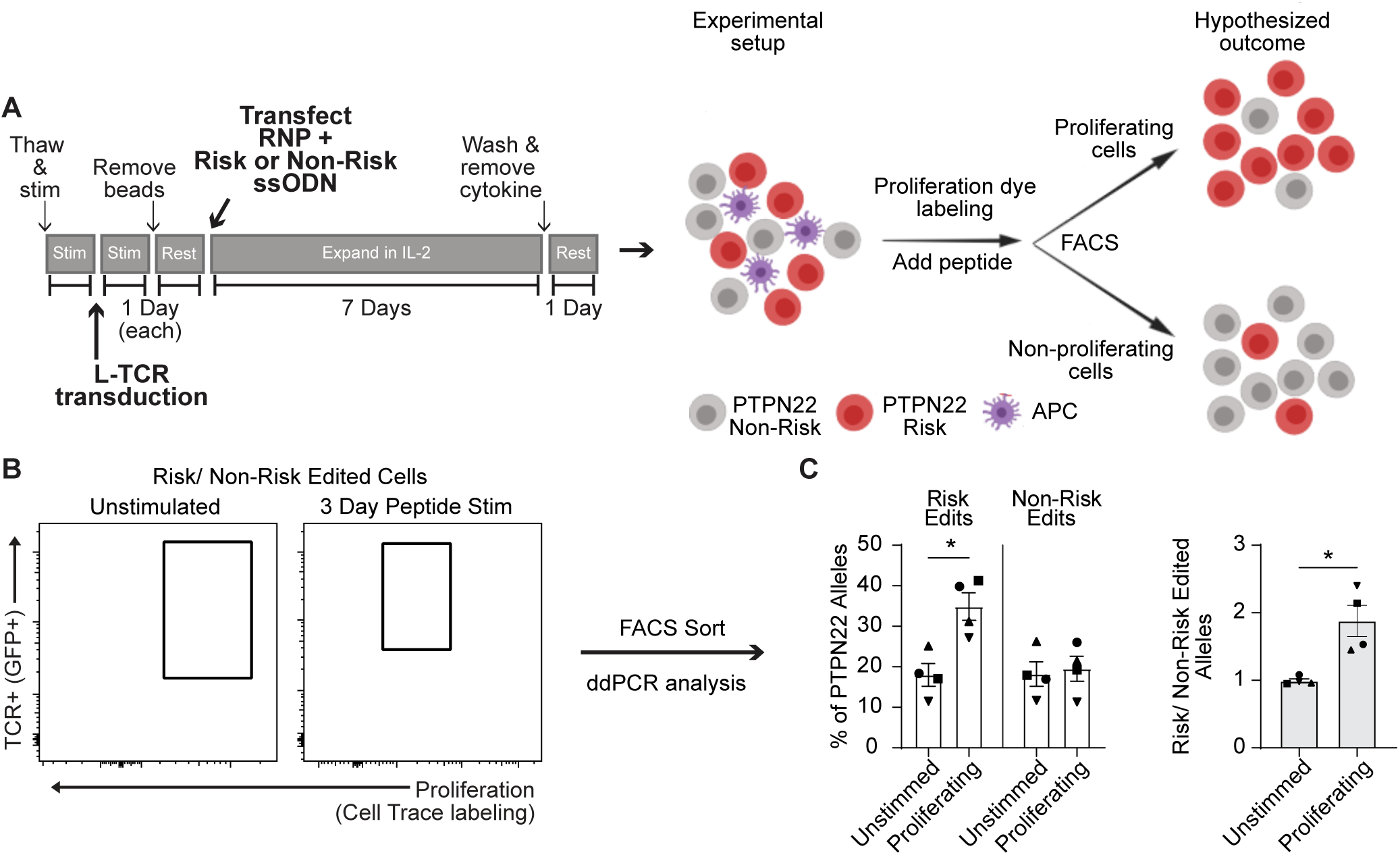
PTPN22 risk edited T cells show increased proliferation compared to non-risk edited cells under L-TCR stimulation. Cord blood CD4 T cells from human donors were transduced with L-TCR lentivirus prior to gene editing into 2 populations (as described in Figure 1): *PTPN22* risk edited or non-risk edited, then stimulated with cognate peptide. (A) Workflow shown in figure 3, depicting the use of *PTPN22* SNP locus editing DNA repair templates and an experimental model describing how the *PTPN22* R620W coding sequence’s impact on proliferation was predicted and would be assessed. (B) A representative FACS plot showing gating of L-TCR+ populations before peptide stimulation and 3 days post peptide stim. (C) ddPCR analysis of edited *PTPN22* alleles in FACS sorted populations. n=4 independent human donors (shapes correspond to individual donors). All summary data analyzed by paired T test. * p<0.05.

## Discussion

In this paper we have utilized a robust gene editing platform in primary human cord blood CD4 T cells to efficiently alter *PTPN22* non-risk primary human T cells to express the *PTPN22* R620W risk variant as well as relevant control non-risk or knockout edited cell populations. This method of gene editing should be applicable to many cell types and could be broadly applied to study the impact of candidate disease associated alleles (identified by GWAS or other sequencing strategies) within primary human cell populations. Notably, our data clearly demonstrate that *PTPN22* risk variant expressing T cells exhibit a phenotype similar to PTPN22 KO T cells, leading to increased TCR induced activation responses. These findings support the conclusion that the *PTPN22* risk variant leads to a loss in negative TCR regulatory function. Further, using transgenic TCR delivery and cognate peptide mediated activation, in tandem with our gene editing platform, we show that PTPN22 deficiency predominantly impacts T cells activated in response to low avidity TCR engagement. In contrast, PTPN22 function is largely dispensable in T cells receiving a high avidity TCR signal. Together, this data suggests that the *PTPN22* risk variant may contribute to autoimmune responses, at least in part, by selectively expanding the pool of lower avidity, self-reactive T cells and skewing such cells toward a pro-inflammatory phenotype.

Our editing platform utilizes a designer nuclease that generates a dsDNA break in close proximity to the target site for HDR editing. This combined approach allowed for: a) generation of edited cell populations with high rates of HDR; b) equivalent HDR rates for alterative coding changes within the target gene via use of a series of ssODN donors; and c) achieving these editing outcomes while controlling for the overall HDR/NHEJ ratio. An additional strength of this approach is that the location and small footprint of these edits allows for physiological levels of expression of edited PTPN22 through use of the endogenous promoter and 3’ UTR, and editing in this location is also unlikely to disrupt the formation of natural PTPN22 splice variants (45). Of note, *PTPN22* risk and non-risk ssODNs, while producing identical HDR rates, demonstrated reduced HDR relative to editing using the PTPN22 KO ssODN. This modest difference likely reflects the requirement to replace endogenous nucleotides immediately 5’ and 3’ to the DNA cut site to achieve risk vs. non-risk edits, while the KO ssODN only required insertion of new nucleotides. These minor differences aside, our editing platform should be readily adaptable for study of targeted edits across the genome in future studies of human primary cell populations.

We demonstrate that the *PTPN22* risk variant can drive expansion of primary human T cells expressing a low avidity TCR, findings that mimic PTPN22 KO T cells. Furthermore, we show that loss of PTPN22 activity promotes the development of a pro-inflammatory phenotype in T cells using a low avidity TCR. These combined findings are consistent with in vivo functional impacts observed in human and mouse studies. Indeed, while previous human data and mouse models have led to contradictory conclusions regarding the impact of the risk variant on proximal TCR signaling, both humans and mice with the *PTPN22* risk variant exhibit expanded memory T cell and Th1 populations (22, 25, 27, 32). However, while our editing design provides a useful means to address questions related to TCR engagement, this approach has limitations in addressing developmental outcomes. For example, while mouse models suggest that the *PTPN22* risk variant alters thymic selection (27) others have shown that PTPN22 KO has no impact on the TCR repertories (29, 31, 32, 51). As this study shows the risk variant functions similarly in human T cells in comparison with published murine models, it suggests that returning to murine models to investigate specific contexts where PTPN22 might impact selection and tolerance are likely to be informative.

Of note, in this study we explored the impacts of the *PTPN22* risk variant and knockout specifically in cord blood CD4 T cells to minimize environmental impact between T cell donors. Although previous work has revealed similar findings for PTPN22 KO in adult T cells (30), it remains possible that similar risk variant editing experiments using adult peripheral blood may yield different results. Also, CD4 T cells were chosen for this study based on well-documented use in gene editing models and likely key roles in PTPN22 associated autoimmune diseases (13). Importantly, PTPN22 is expressed in multiple immune and hematopoietic subsets and has been shown to have multiple functions (11, 12, 18). Therefore, our dataset informs on a single signaling program and PTPN22 may contribute to autoimmune disease via other pathways.

While T cells expressing a lower avidity TCR were primarily impacted in our studies, we cannot eliminate potential alternative impacts of the TCRs studied in our models. The TCRs evaluated exhibited distinct differences in both EC_50_ (43) and triggered proliferative response to an identical IGRP derived peptide (Fig. 3, C) (43). However, while these data points support a working hypothesis of TCR avidity determining the functional impact, we cannot rule out the role for possible differences in TCR disassociation rates and/or overall duration of TCR engagement. Further, while our approach provides a robust system to assess the impact of the risk variant in primary human T cells, our findings are derived from a comparison of only two specific TCRs. The identification of similar pairs of “higher” and “lower” avidity TCRs recognizing identical self-peptides will be useful in assessing our model and its interpretation across a broader pool of TCR interactions.

Through gene editing we demonstrate that *PTPN22* risk variant expression in human T cells mediates a partial loss of function phenotype that mirrors findings derived from mouse models. These data, however, do not explain the marked differences in T cell responses described in mouse models vs. natural carriers of the *PTPN22* risk variant. Natural carriers of the risk variant exhibit hyporesponsive T cells, with reduced TCR induced activation, and reduced cytokine responses (22, 23). In previous work, we showed that loss of function in another key negative regularity pathway in T cells leads to overexpression of functionally related proteins to maintain homeostasis. For example, knockout of the IL-2 regulatory gene, *PTPN2,* in primary human T cells results in a transient increase in IL-2 signaling, but expansion of PTPN2 KO cells in high cytokine media leads to reduced IL-2 responses due to overexpression of the regulatory protein SOCS3 (30). Similar compensatory events may be operable in T cells in carriers of the *PTPN22* risk variant, where reduced proximal negative regulatory events might lead to expression of alternative regulatory proteins. Recent work using optogenically controlled TCR signaling has shown that T cell activation is dynamic, requiring optimal frequencies of TCR engagement relative to cessation of signal to mediate CD69 expression and subsequent activation; however, events that limit T cell responses may be more static, with expression of TIM3, for example, increasing regardless of signaling frequency (52). As loss of PTPN22 function specifically augments low avidity TCR signaling, it is possible tonic TCR signaling is also impacted and may drive additional events that function to limit TCR signals in carriers of the risk variant over time. Finally, it is important to highlight the environmental differences between murine models and primary human T cell donors. As opportunity for exposure to antigen increases, this may drive T cells from human carriers towards a reduced responsiveness to TCR engagement via impacts that are not operative in SPF murine models.

In summary, the data presented within this study supports a model where the *PTPN22* R620W risk variant impacts human T cell function via a reduction in negative regulation of TCR signaling; findings similar to murine models, but in contrast to the T cell signaling phenotype in carriers with the risk variant. In association with previous human studies, our findings suggest that the impact of the risk variant on T cell function is likely dynamic, leading to the generation of populations of hyperresponsive T cells, but also allowing for secondary impacts on T cell gene expression that may lead to hyporesponsive T cells over time or after environmental exposures. This would help explain how PTPN22 could influence risk of diverse autoimmune diseases such as type 1 diabetes and rheumatoid arthritis, diseases with distinct ages of onset and tissue targets. Taken together, these observations suggest that PTPN22 contributes to autoimmunity by establishing temporal windows of risk for specific diseases that coincide with periods of altered T cell responsiveness.

## Acknowledgements

We thank Rich James (Seattle Children’s Research Institute, Seattle, WA) for his helpful comments and insight.

## Author contributions

WA, JHB, and DJR conceived the study. WA and DJR developed the experimental designs. WA performed experiments and analyzed data with input from DJR. FW, KC and PSL identified and generated the DNA constructs for the L-TCR and H-TCR lentiviruses. WA and DJR wrote the manuscript. All authors reviewed and approved the manuscript.

---

**Supplemental Figure 1.**
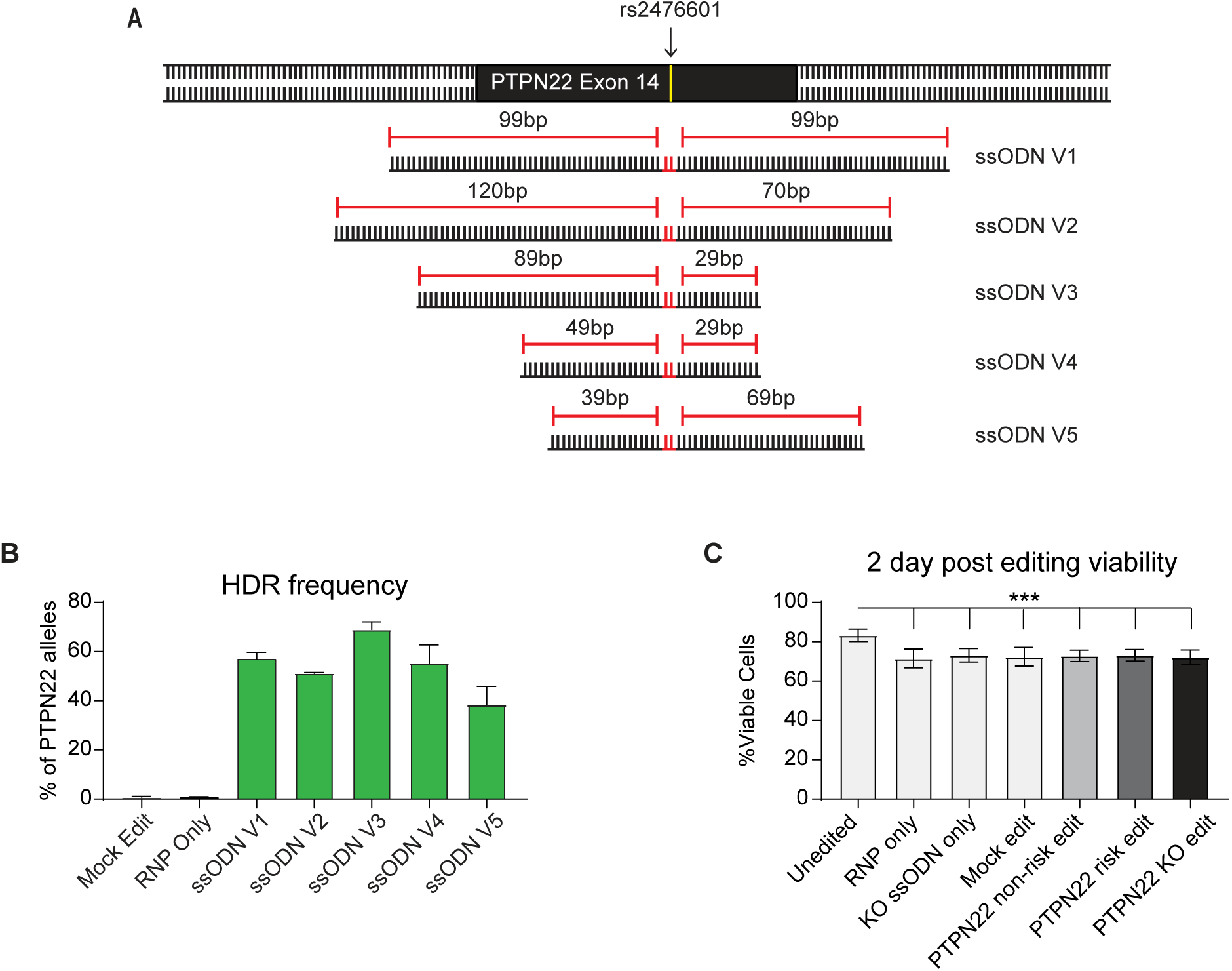
Gene editing efficiency of *PTPN22* SNP variants is impacted by donor DNA template design. (A) ssODN repair template designs tested to introduce *PTPN22* risk SNP at rs2476601 site in non-risk T cells. Cells were electroporated with a *PTPN22* targeting Crispr/Cas9 RNP (Figure 1A), and one of 5 ssODN variants, all introducing a 2bp edit resulting in producing the R620W variant and blocking further binding of the Crispr/Cas9 RNP. All ssODN variants are complementary to the non-target strand of the RNP. (B) Average HDR rates resulting in the *PTPN22* risk variant achieved with ssODN variants. CD4 T cells were edited as in Figure 1B and processed for gDNA 2 days post editing. Rates of HDR were determined by ddPCR analysis (n = 2-4 independent donors, and 1-2 independent experiments). For ssODN design optimization CD4 T cells were derived from human adult PBMCs. (C) After editing with optimized ssODN templates, cord blood CD4 T cells were expanded in IL-2 for 7 days. At 2 days post editing, cells were stained with viability dye and assessed via flow cytometry. Percent viable reflects the percent of events collected from each culture that were identified as single, live, lymphocytes. n=4, independent human donors. For summary graphs lines and error bars represent mean +/- SEM, no datapoints are from replicate human donors. Analyzed by matched one-way ANOVA with Tukey’s correction. RNP - ribonucleoprotein, ssODN - single stranded oligo-deoxynucleotide, NHEJ - non-homologous end joining, HDR - homology directed repair. *** p<0.001.

**Supplemental Figure 2.**
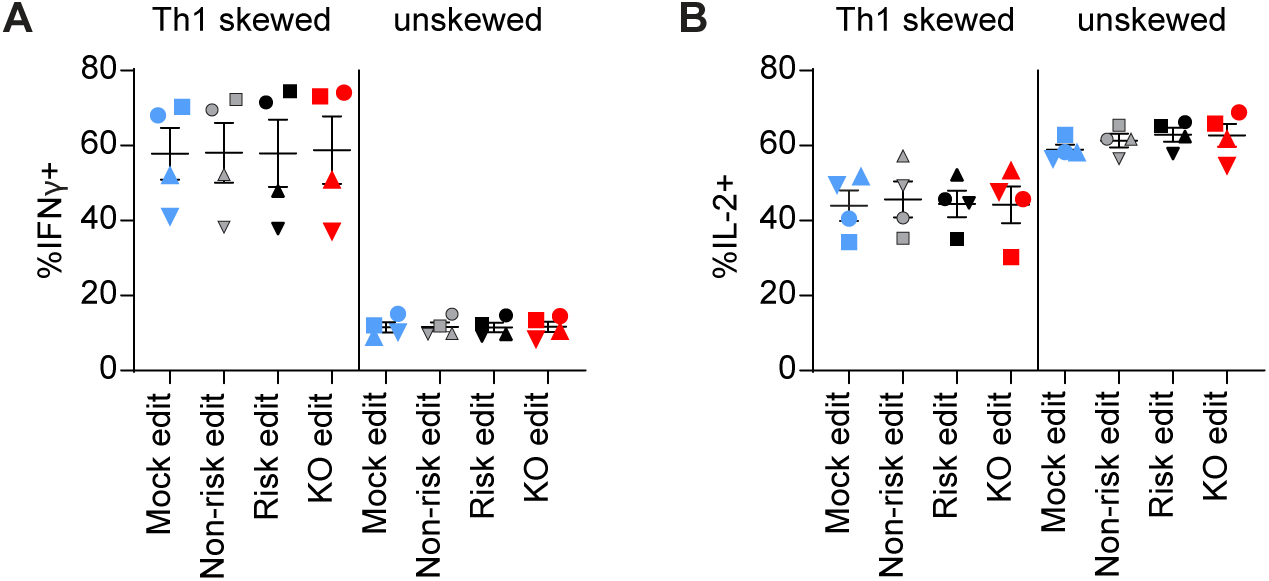
*PTPN22* editing in T cells does not impact IL-12 induced Th1 skewing. Cord blood CD4 T cells from human donors were edited as described in Figure 1, then stimulated with plate bound anti-CD3 for 72 in either Th1 skewing media (IL-12, anti-IL-4, IL-2, and soluble anti-CD28) or non-skewing media (IL-2 and soluble anti-CD28). Cells were then stimulated with PMA and ionomycin with monensin for 5 hours, then fixed and stained for flow cytometry assessment of cytokine production of (A) IFNγ and (B) IL-2. n=4 independent donors (shapes correspond to individual donors), lines and error bars represent mean +/- SEM. All data is from 2 independent experiments.

**Supplemental Figure 3.**
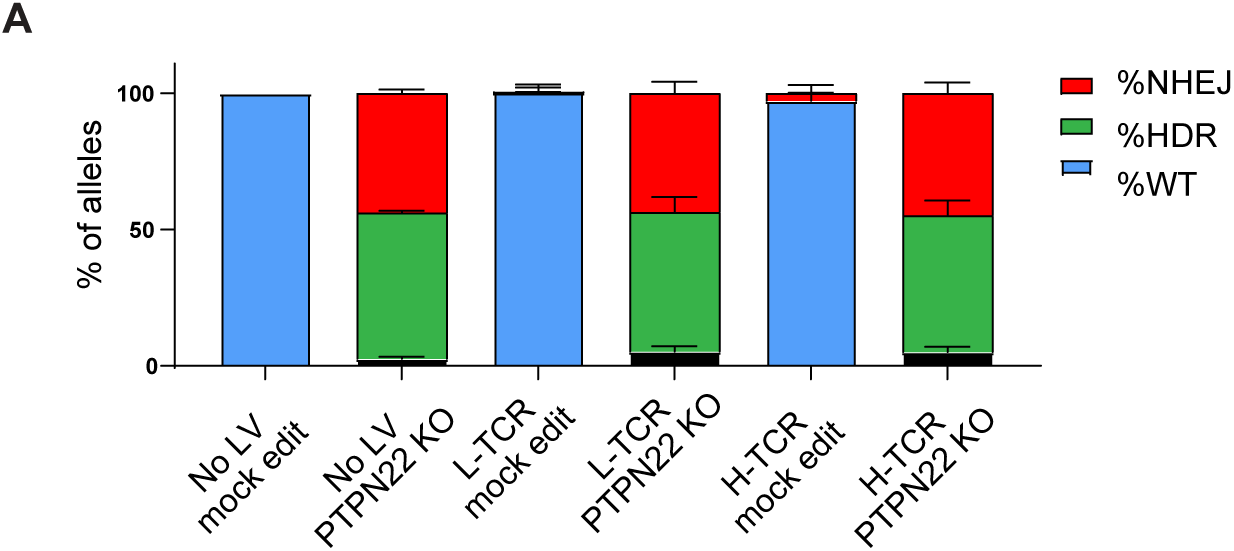
*PTPN22* gene editing in cord blood CD4 T cells is not impacted by lentiviral transduction. **(A)** ddPCR analysis of *PTPN22* editing in human cord blood CD4 T cells using a KO ssODN repair template (Figure 1A) after lentiviral (LV) delivery of antigen specific TCRs (Figure 3A). n=4-6 independent donors, columns and error bars represent mean +/- SEM. Figure reflects data compiled across 5 independent experiments, with each column representing data from 2 to 3 independent experiments.

**Supplemental Figure 4.**
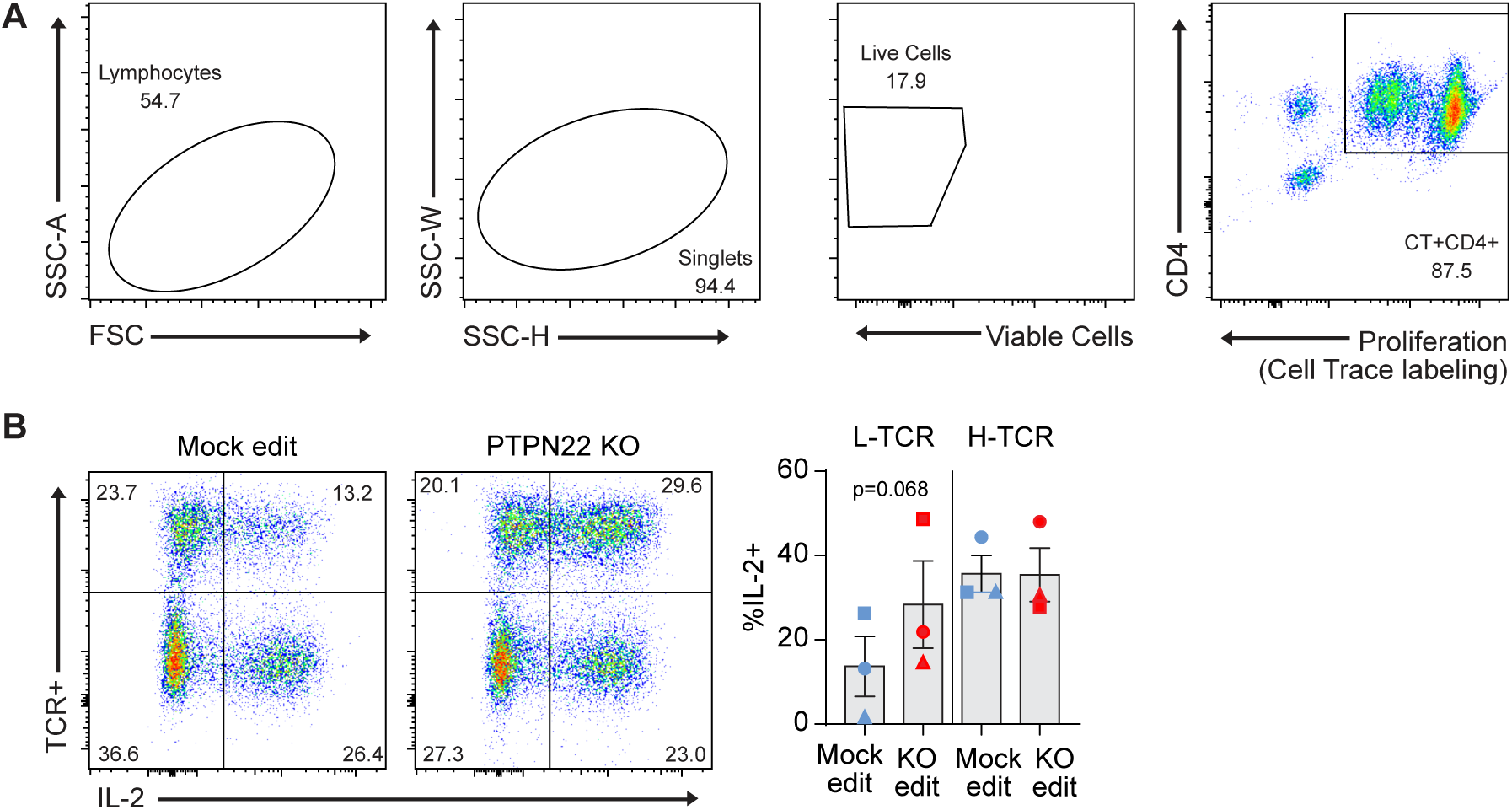
*PTPN22* KO cord blood CD4 T cells show trend toward increased IL-2 secretion under L-TCR stimulation. Cord blood CD4 T cells from human donors were edited as described in Figure 3, then stimulated with peptide loaded APCs for 72 hours. (A) Flow cytometry gating strategy used to identify antigen specific TCR+ cells for cytokine production analysis. (B) Representative flow plots of IL-2 expression in mock edited or PTPN22 KO cells transduced with L-TCR and summary data of %IL-2 secreting *PTPN22* edited/TCR+ cells. n=3 independent human donors, from 2 independent experiments). All summary data analyzed by paired T test. Columns and error bars represent mean +/- SEM, and shapes correspond to individual human donors.

**Supplemental Figure 5.**
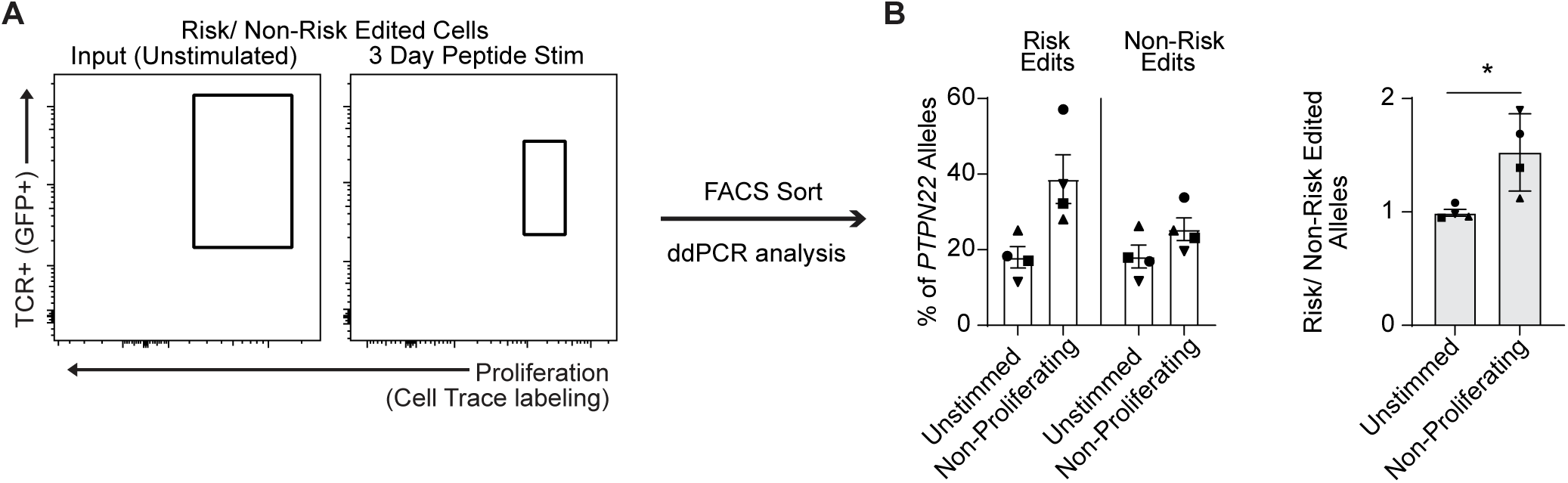
Proportion of *PTPN22* risk edited T cells increases compared to non-risk edited cells under L-TCR stimulation even in non-proliferating cells. Cord blood CD4 T cells from human donors were transduced with L- TCR lentivirus prior to gene editing as described in Figure 1 and Figure 6 and mixed in equal proportions as shown in Figure 6A, then stimulated with cognate peptide. (**A**) Representative FACS plot showing gating of L-TCR+ populations before peptide stimulation and 3 days post peptide stim to sort non-proliferating events. (**B**) ddPCR analysis of edited *PTPN22* alleles in FACS sorted populations. n=4 independent human donors (shapes correspond to individual donors). All summary data analyzed by paired T test. * p<0.05.

